# Shifting up a gear with iDNA: from mammal detection events to standardized surveys

**DOI:** 10.1101/449165

**Authors:** Jesse F. Abrams, Lisa Hörig, Robert Brozovic, Jan Axtner, Alex Crampton-Platt, Azlan Mohamed, Seth T. Wong, Rahel Sollmann, Douglas W. Yu, Andreas Wilting

## Abstract

1. Invertebrate-derived DNA (iDNA), in combination with high throughput sequencing, has been proposed as a cost-efficient and powerful tool to survey vertebrate species. Previous studies, however, have only provided evidence that vertebrates can be detected using iDNA, but have not taken the next step of placing these detection events within a statistical framework that allows for robust biodiversity assessments.
2. Here, we compare concurrent iDNA and camera-trap surveys. Leeches were repeatedly collected in close vicinity to 64 camera-trap stations in Sabah, Malaysian Borneo. We analyze iDNA-derived mammalian detection events in a modern occupancy model that accounts for imperfect detection and compare the results with those from occupancy models parameterized with camera-trap-derived detection events. We also combine leech-iDNA and camera-trap data in a single occupancy model.
3. We found consistent estimates of occupancy probabilities produced by our camera-trap and leech datasets. This indicates that the metabarcoding of leech-iDNA method provides reasonable estimates of occupancy and can be a suitable method for studying and monitoring mammal species in tropical rainforests. However, we also show that a more extensive collection of leeches would be needed to assess mammal biodiversity with a similar robustness as with camera traps. As certain taxa were only detected in leeches, we see great potential in complementing camera-trap studies with the iDNA approach, as long as the collection of leeches follows a similar robust and standardized sampling scheme.
4. *Synthesis and applications*. The approach we describe here is not restricted to the processing of leech samples, but can be used for the analysis of other iDNA and environmental DNA (eDNA) data. Our study is the first step to shift the application of e/iDNA studies from opportunistic ad-hoc collections to systematic surveys required for long-term wildlife populations and biodiversity monitoring programs.

## 1. Introduction

To halt further biodiversity loss, parties of the Convention of Biological Diversity (CBD) have agreed to track and report progress towards the Aichi Biodiversity targets and to the Intergovernmental Science-Policy Platform on Biodiversity and Ecosystem Services (IPBES). To do so, rigorous monitoring of wildlife, using fast and efficient tools, is necessary (Bush *et al.*, 2017). The protocol for such assessments already exists: gather detection events, analyze these using modern statistical models, and track population status over time (Bush *et al.*, 2017). However, detecting species, particularly in tropical rainforests, remains a challenge as species are often secretive and occur in remote areas.

Today, numerous methods are used to gather detections of mammals, all of which are time- and labor-intensive. Camera trapping has proven to be the most labor-efficient method (Roberts, 2011) and now plays an important role in wildlife management, allowing researchers to record a wide range of species in remote terrain over long time periods (Abrams *et al.*, 2018; Burton *et al.*, 2015; Trolliet *et al.*, 2014). With the increased use of camera-trapping surveys, the methods for processing and statistically analyzing the data have also advanced (Burton *et al.*, 2015; MacKenzie *et al.*, 2006). However, the use of camera traps remains limited by difficult setup and high capital and maintenance costs.

An alternative or complement is environmental DNA (eDNA), which refers to the DNA that can be collected from a variety of environmental samples such as soil, water or faeces (Bohmann *et al.*, 2014; Thomsen & Willerslev, 2015; Ilshige *et al.*, 2017). Recent methodological advances, namely amplicon-based high-throughput sequencing or ‘metabarcoding’, now also allow the reliable reading of such DNA sources (Abrego *et al.*, 2018). Invertebrate-derived DNA (iDNA) is an offshoot of eDNA, where terrestrial vertebrates can be detected via their DNA that was ingested by invertebrates (Schnell *et al.*, 2012, 2015; Calvignac-Spencer *et al.*, 2013a, 2013b; Tessler *et al.*, 2018; Weiskopf *et al.*, 2018). Sanguivorous species such as leeches (Schnell *et al.*, 2012, 2018), mosquitos (Kent and Norris, 2005), or ticks (Gariepy *et al.*, 2012) are commonly used, and invertebrates that feed on vertebrate fecal matter or carcasses, such as dung beetles, blow flies, and carrion flies have also been employed (Calvignac-Spencer *et al.*, 2013a, 2013b; Lee, Sing, and Wilson, 2015; Rodgers *et al.*, 2017; Schubert *et al.*, 2015; Somervuo *et al.*, 2017). Although these initial studies provide proof of principle that vertebrates can be detected using iDNA, they have been restricted to opportunistic collections of invertebrates and only been used to compile species lists. The sampling and the analyses have not been carried out in ways that allow statistically robust assessments of species or community trends and species population status.

The issue is that e/iDNA, like all detection methods, is imperfect: the non-detection of a species by e/iDNA in a location does not prove the absence of that species in that location. Accounting for imperfect species detection requires a well-designed sampling scheme, combined with a statistical method known as occupancy modelling (MacKenzie *et al.*, 2002) to estimate the true spatial extent of species presence from detection events. Occupancy models, which estimate the percentage of area occupied and the probability of occupancy at any given site, have been widely used on camera-trap data (Burton *et al.*, 2015), but have so far only been proposed for e/iDNA data (Schnell *et al.*, 2015). Occupancy modelling (Fig. S1) uses detection/non-detection data collected from repeated sampling of the same locations. Under the assumption that the target vertebrate community does not change between sampling events, known as the ‘closed-population’ assumption, the repeated collection of species detection/non-detection data can be used to estimate the probability of species occurrence correcting for detection probability <1. Furthermore, both detection and occupancy probability can be modelled as functions of covariates.

**Figure 1.**
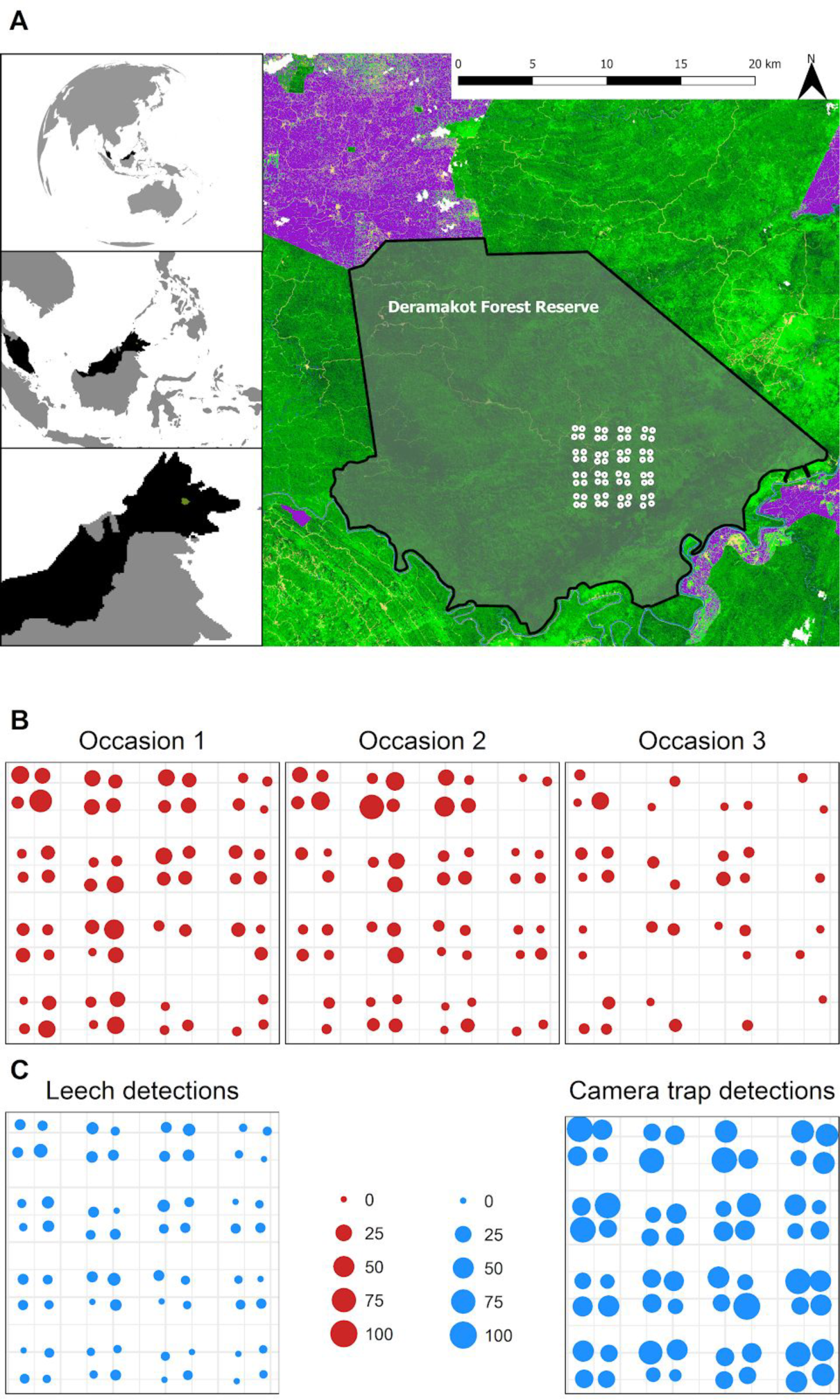
(A) The study site, Deramakot Forest Reserve, in Malaysian Borneo, with the locations of the 64 camera-trap and leech sampling stations (white circles). (B) The number of leeches sampled (indicated by the size of the circles) in Deramakot on the three sampling occasions. (C) The number of detections for camera-trap and leech surveys (represented by the size of the circles).

Here, we present an approach that makes this much-needed shift from proof-of-principle studies using iDNA to gather detection events to using iDNA as an input to statistically robust biodiversity assessment. We carried out standardized and repeated collections of terrestrial haematophagous leeches in Sabah, Malaysian Borneo, and we employed a molecular pipeline (Axtner *et al.*, 2018) that minimizes the risk of false-positive species detections. We compared leech-derived and camera-trap species richness estimates and investigated the detection bias towards smaller or larger species. We then analyzed the leech-derived species detections within an occupancy-modelling framework that accounts for imperfect detection and compared the results to estimates of detection and occupancy probability from concurrently collected camera-trap data. Finally, we combined the leech iDNA and camera-trap data in a single occupancy model to evaluate the opportunities of combining e/iDNA data with conventionally collected biodiversity data.

## 2. Methods

### 2.1 Study area and data collection

We conducted camera-trap surveys and leech collection in the Deramakot Forest Reserve in Sabah, Malaysian Borneo (Fig. 1), which covers an area of approximately 55,500 hectares (ha) of mixed dipterocarp forest. A systematic camera-trapping survey was carried out from February - May 2015. Camera traps were deployed in a clustered design with each cluster consisting of a square of four camera-trap stations, spaced 500 m apart (Fig. 1). 16 clusters were established in a 4 × 4 formation with a distance of 1.5 km between cluster centers. Each station consisted of two Reconyx PC850 white-flash camera traps facing each other, operating 24 hours/day, and left in the forest for a minimum of 60 days (for details, see Abrams *et al.*, 2018). Two type of leeches, tiger and brown leeches, were collected concurrent to camera trapping. As taxonomy within the genus *Haemadipsa* is currently not resolved (see Schnell *et al*., 2015), we refer to the types only, although tiger leeches are described as *Haemadipsa picta* and brown as *Haemadipsa zeylanica* (Fogden & Proctor, 1985). Tiger leeches, the larger of the two types, live in small trees and bushes, while the smaller brown leeches occur mainly on the ground. This difference in behavior may lead to different preferences in host species (Schnell *et al.*, 2015). Samples were taken within a 20 x 20 m sampling plot around the camera-trap stations. Sampling was repeated three times with approximately 30 days between sampling instances (at setup, check, and collection of camera traps). The leeches were immediately placed in RNAlater and stored at −20 °C. All leeches of the same type (tiger or brown) from the same site and occasion were pooled and processed as one sample.

### 2.2 Laboratory procedures and taxonomic assignment

We implemented a novel e/iDNA workflow to extract raw species detections from leech iDNA (see Axtner *et al*. (2018) for a full description of our methods and Fig. S2). In short, leech samples were first digested, and each sample was split into two extraction replicates, from which DNA extraction was carried out. Of each extraction replicate, we PCR-amplified three vertebrate mitochondrial markers twice, *12S*, *16S* and *cytochrome-b* twice. This resulted in 12 PCR replicates for each leech sample (2 technical replicates × 2 PCR replicates × 3 markers). We used a two-step, twin-tagging PCR strategy to produce double-labeled PCR libraries, which allowed us to implement a high throughput workflow and to minimize the risk of sample misidentification (Axtner *et al.*, 2018). PCR products were sequenced using Illumina MiSeq, and after sample demultiplexing and processing, we assigned each haplotype to a taxonomy using a curated reference database (Axtner *et al*., 2018) and the PROTAX software (Somervuo *et al.*, 2016).

**Figure 2.**
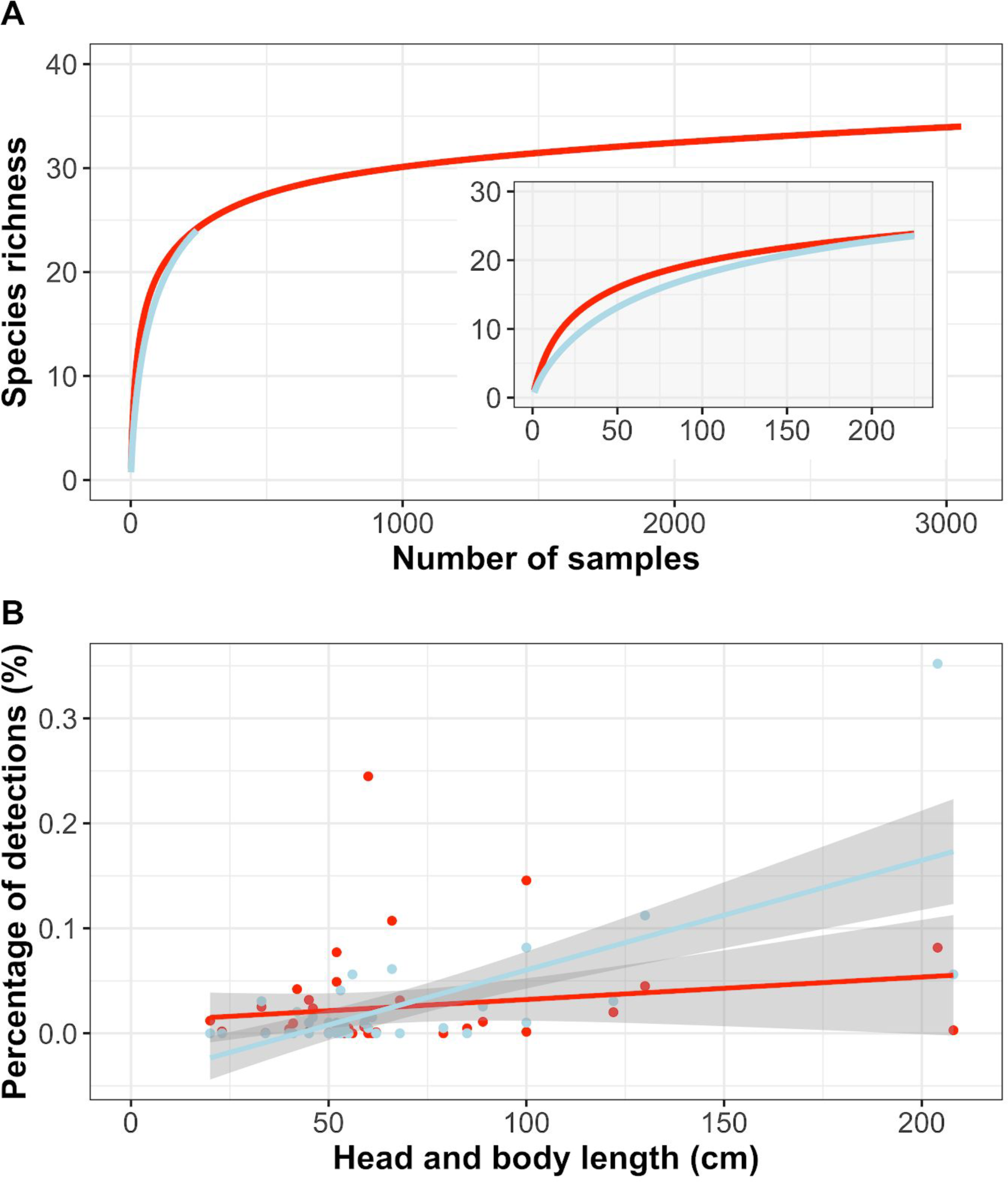
(A) Species-accumulation curves constructed for camera-trap detections (red line) and leech detections (light blue). The main plot shows the mammal species-accumulation up to the total of 3055 camera-trap detections while the inset shows the accumulation up to the first 250 detections. (B) Correlation between detections and the average head and body length of species. The solid lines represent the best fit line, the grey shaded areas represent the 95% confidence intervals.

We followed the *lax* criteria by Axtner *et al*. (2018) for accepting an assignment, in which a species detection was accepted if it appeared in at least two PCR replicates.

### 2.3 Data analysis

For the leech and camera-trap surveys, we computed mammal species accumulation curves in R using the function *specaccum* from the *vegan* package (Oksanen *et al.*, 2007). Earlier leech studies proposed that leeches might be better suited for the detection of smaller bodied species than are camera traps (Weiskopf *et al.*, 2018). We investigated whether detections in brown or tiger leeches were associated with host’s body length by checking for correlation between the percentage of detections of a given species and the species body length in both tiger and brown datasets using Spearman rank. Species head and body length data was taken from the Pantheria database (Jones *et al.*, 2009).

For the occupancy analysis, we used a subset of 11 species that were detected multiple times in both the leech and camera-trapping surveys. We adopted the hierarchical formulation of occupancy models by Royle & Dorazio (2008) and used single-species, single-season models (MacKenzie *et al.*, 2002). We defined a total of six sampling occasions for the occupancy analysis of the leech data, based on the two types of leeches and the three sampling events. We excluded all stations in which no mammals were detected in any of the six leech samples (i.e. stations with no leech-derived mammal data). This resulted in subsets of 49 stations for the occupancy analysis, for which we created detection histories.

We observed a difference in raw detection rates between tiger and brown leeches (see Fig. S3 and Table S1), so we included a categorical covariate on detection for the two leech types. The probability of detecting a mammal species in a leech DNA sample likely also depends on the number of leeches collected within this sample, as well as the number of other species detected in the sample, as species with low DNA amounts in the sample might not amplify in the PCR, if more abundant DNA of other species is present. Therefore, we included both the number of leeches per sample (we referred to this as *effort^leech^*), as well as the number of species detected per sample (referred as *detection^leech^*) as covariates on detection probability.

**Figure 3.**
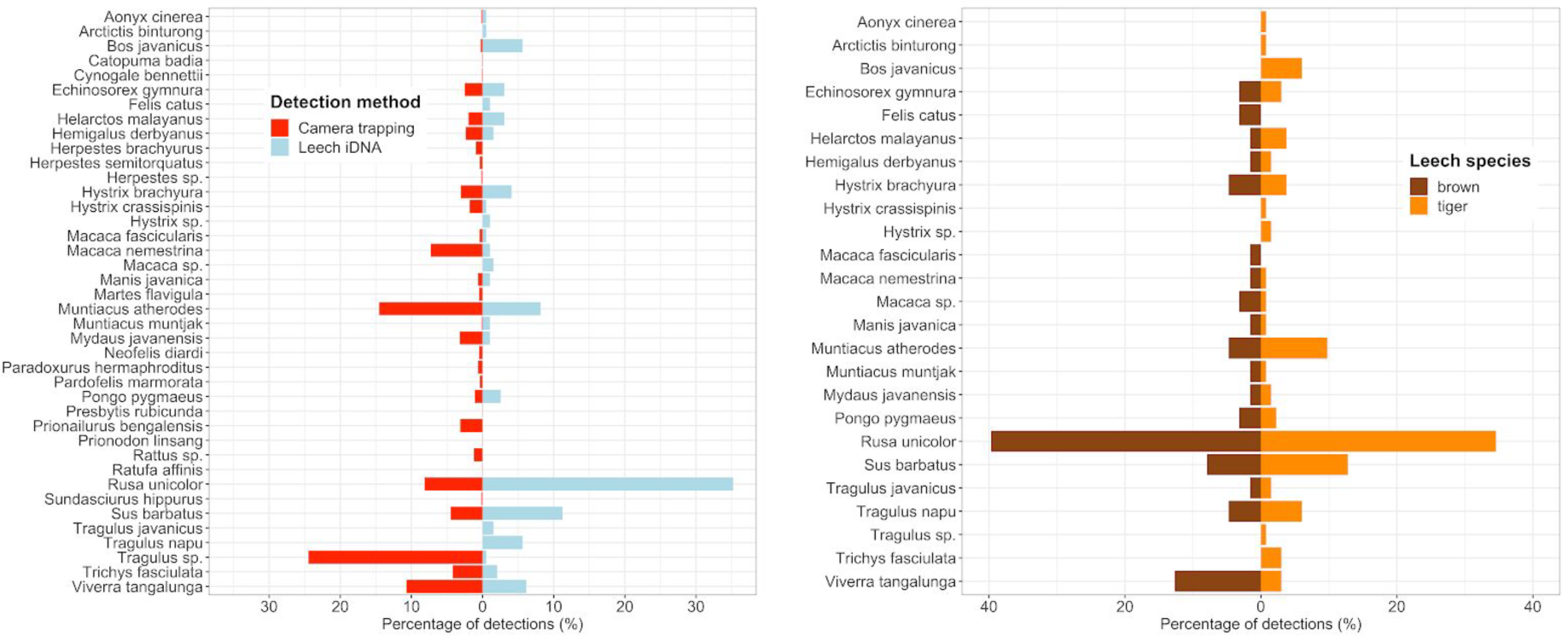
(A) Mammal species compositions of photographs from 1,334 camera-trap nights which generated 2,733 detections (left, red bars) and from 242 pooled leech samples, which generated 196 detections (right, light blue bars). (B) Percentage of successful detections of species in 116 pooled tiger leech samples, which generated 133 detections (right, orange bars) and 126 pooled brown leech samples which generated 63 detections (left, brown bars).

To compare the detection probabilities (p) and the occupancy probabilities (ψ) of the leech dataset (ψ_leech_, p_leech_) with the camera-trapping dataset (ψ_ct_, p_ct_), we prepared the camera-trap data for the same subset of species and stations for occupancy analysis using the R package *camtrapR* (Niedballa *et al.*, 2016) with an occasion length of seven days. Since not all occasions were sampled for the full seven days (e.g. due to camera-trap failure) we accounted for this *effort* on the detection probability. Both effort and the number of species detected per sample are indexed by site (*j*) and occasion (*k*) (for a full model description see the Supporting Information).

We implemented the models in a Bayesian framework using JAGS (Plummer, 2003) accessed through R version 3.4.2 (R Core Team, 2017) using the package *jagsUI* (Kellner, 2015). We report results as posterior mean (in cases of skewed posterior distributions, the mode) and 95% Bayesian credible intervals (95BCI, the 2.5% and 97.5% percentiles of the posterior distribution). We evaluated the performance and accuracy of the single species occupancy models from both the leech-iDNA and camera traps based on the 95BCI of detection and occupancy probabilities.

Last, we evaluated the value of combining camera-trapping and leech detections in a joint analysis. In the combined models, we included a categorical covariate for the detection method so that we could estimate the detection probabilities for the camera traps, brown leeches, and tiger leeches independently, but draw on all sources to estimate occupancy.

Our survey design with the 500 - 1000 m spacing between camera-trap stations will most likely lead to spatial autocorrelation for some species, creating some bias in occupancy estimates (Legendre, 1993; Dormann, 2007). However, since the data were collected according to the same survey design, the spatial autocorrelation bias will likely be the same for leech and camera-trap surveys, and so any bias will not affect our comparison of camera traps and iDNA.

## 3 Results

A total of 1,532 leeches (801 brown; 731 tiger) were collected during the survey, with the number of leeches sampled varying between stations and sampling occasions (Fig. 1). Leeches of both types were not detected at every site on every occasion. The number of leeches collected decreased from 762 to 576 to only 194 in the first, second, and third sampling occasions, respectively. Leeches of the same collecting occasion and type were pooled for a total number of 126 brown-leech and 116 tiger-leech samples (i.e. a collection tube). From these 242 samples, 196 mammal detection events were made after sequencing and bioinformatic processing. In 1,334 camera-trapping nights we obtained 3,055 independent records from camera traps, resulting in 2,733 mammal detection events. The camera-trap data had 31 identifiable mammal species, while the leech data had 22 mammal species (Table 1, Fig. 2A). All mammal species detected via leech iDNA are known to occur in the study area, and two of the species, *Arctictis binturong* and *Felis catus* were not detected by the camera traps. Additionally, using the leech iDNA, we were able to distinguish two species of mouse deer, *Tragulus javanicus* and *Tragulus napu*, which was difficult and often impossible from the camera-trap photographs. In some cases we were not able to assign species-level taxonomies to sequences from the leech iDNA, due to an incomplete reference database or a low-confidence assignment from PROTAX (usually caused by low inter-specific sequence diversity). The most frequently identified species in the leech samples was *Rusa unicolor* with 69 detections, followed by *Sus barbatus* (22 detections, Table S1). The most identified species in the camera-trap samples were *Tragulus sp*. (669 detections), *Muntiacus atherodes* (398), and *Viverra tangalunga* (293). The species accumulation curve of the leech detections showed a similar increasing trend as the camera-trap dataset (Fig. 2a), but did not reach their asymptotes, contrary to the camera-based curves.

**Table 1.**
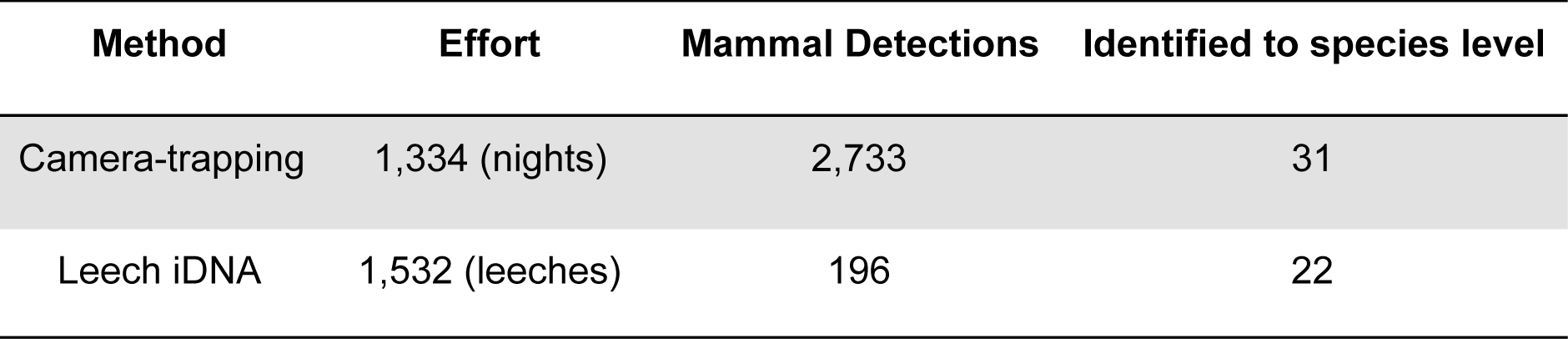
Summary of detections and effort for camera-trapping and leech collection.

A comparison of the detections of mammal species in tiger and brown leeches revealed that the detection rate (detections per samples) in tiger leeches (1.15) was much higher than that of brown leeches (0.5). The smallest mammal we detected was the *Echinosorex gymnura* with a body mass of ~756 gr (Jones *et al.*, 2009), which was also detected in the camera-trap survey. We did, however, record a frog species *Kalophrynus pleurostigma* in the leech dataset. The percentage of detections in the leech data was significantly positively correlated with the head and body length of the species (Fig. 2; Spearman’s rho = 0.491; p-value = 0.0013). This pattern did not extend to the camera-trap data (rho = 0.115, p-value = 0.481, Fig. 2b).

Detection probability (Fig. 4a) varied between species and detection methods. Generally, the estimated detection probability for the camera-trap dataset had smaller CIs. In the leech dataset, estimates of detection probability of species with a low number of detections had very high uncertainty (Fig. 4a). Detection probability in the tiger leeches was higher than in the brown leeches for all but one species, *Viverra tangalunga* (Fig. 4a). The detection probability of brown leeches was lower than that of camera traps for all species, except *Pongo pygmaeus*. Tiger leeches had more success detecting certain species than did camera traps, with a higher estimated detection probability for 5 out of the 11 species. Despite the higher number of detections in the camera-trap dataset, the occupancy models for the two survey methods generated similar occupancy probabilities with overlapping confidence intervals (Fig. 4b). The occupancy estimates from the camera-trap data had narrower BCI for all species except banteng and sambar (Fig. 4b).

**Figure 4.**
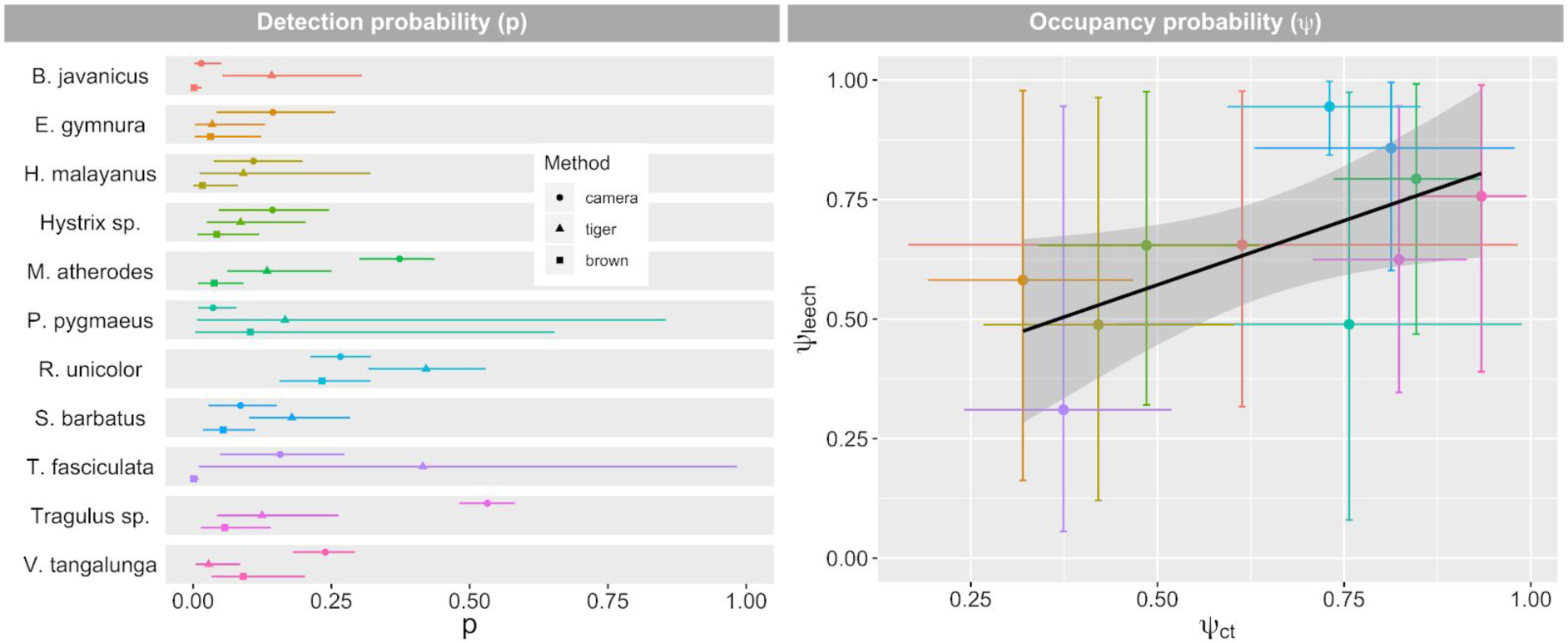
Occupancy and detection probabilities estimated by null single-species occupancy models for independent camera-trap and leech survey data, which include 32 stations. (A) Camera-trap (circles), tiger (triangles), brown (squares) detection probabilities are plotted with their 95% Bayesian CIs. (B) Estimated occupancy probabilities for camera-trap models (x-axis) are plotted against the estimated occupancy probabilities for leech models (y-axis). The vertical and horizontal bars indicate 95% Bayesian CIs for the leech survey and camera-trap survey, respectively. The black line is the best fit line, the grey shaded area represents the 95% confidence interval.

Occupancy estimates for the independently analyzed leech and camera-trap datasets were similar with those from the combined analysis (Figs. 5, S4). When compared to the camera-trap only models, the combined dataset resulted in smaller BCIs in the estimates of occupancy probabilities for the suite of 11 single-species models by an average of 12% (Figs. 5, S4). The suite of 11 species had an average BCI of 0.639, 0.337, and 0.297 for the leech, camera-trap, and combined models, respectively.

**Figure 5.**
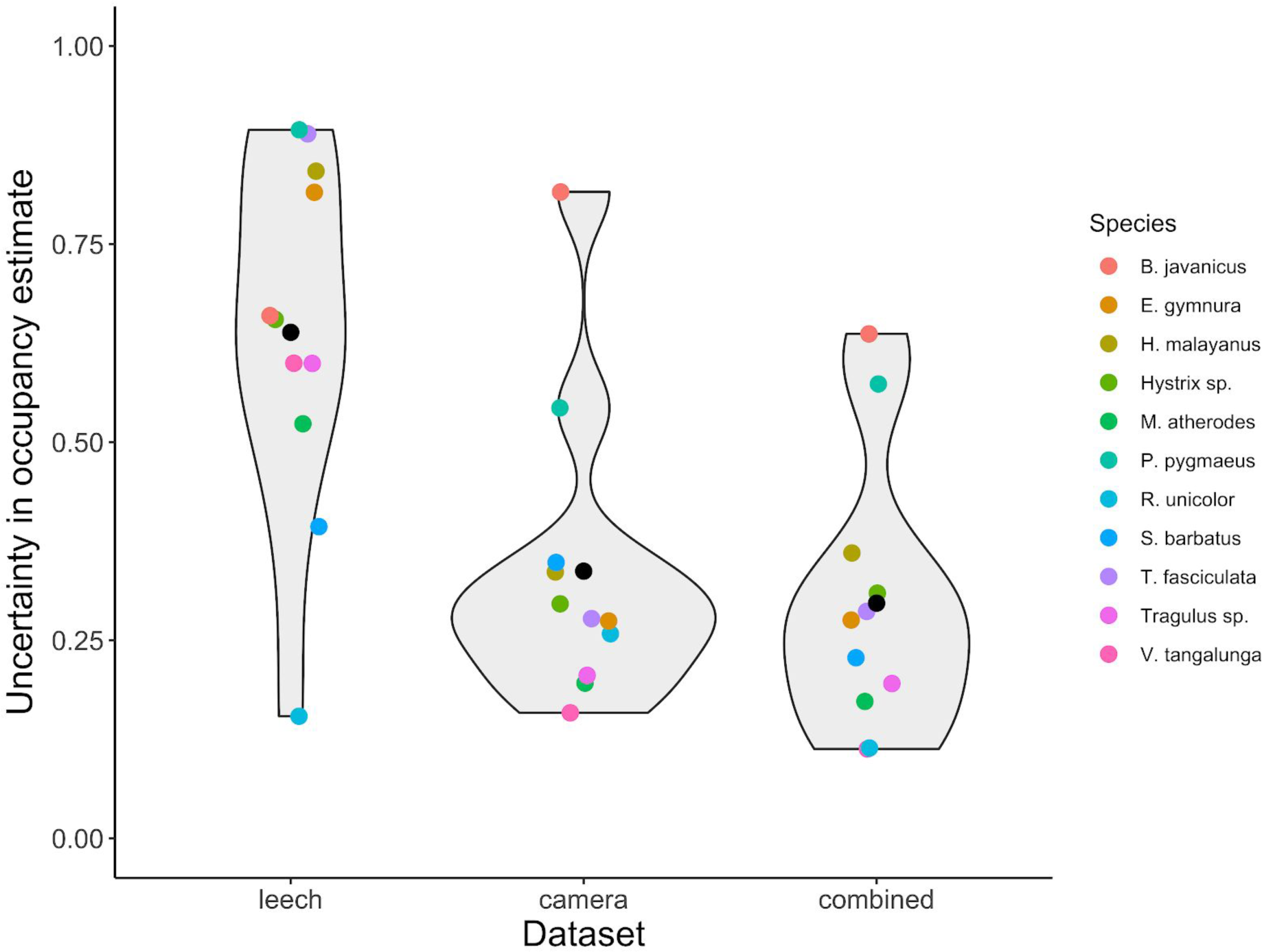
The size of the 95% Bayesian CIs for modeled occupancy probability for 11 species (represented by the colored points) for the leech iDNA, camera-trap, and combined datasets, which includes 32 stations. The black dots for each dataset represent the average size of the occupancy probability CIs. The grey shaded areas represent the distribution and probability density of the size of the confidence intervals estimated from the single species models.

## 4 Discussion and Conclusion

Overall, our results showed that 128 camera traps in 64 stations resulted in more detections and in higher species-richness estimates than did three leech collections at the same stations (Fig. 2A). Although requiring the same amount of field time (three visits), the sampling efforts were very different as the cameras were active for up to 63 nights. Because the initial species-accumulation curves for the camera-trap and leeches were quite similar (Fig. 2A, inset), we expect that increased numbers of leech samples could also achieve the same estimate of total species richness and decreased uncertainty in occupancy estimates, due to higher numbers of repeat samples.

On the other hand, leech sampling success can vary due to season, weather, and microhabitat conditions. Thus, an understanding of the factors that influence leech abundance is an important requirement to adequately design and execute a successful leech study (Schnell *et al.*, 2015). Currently, little is known about the ecology of terrestrial leeches, but earlier studies indicated that they depend on humid conditions and survive the dry season by burying themselves in the soil (Nesemann and Sharma, 2001). In our study, the number of leeches collected varied largely between the stations and decreased with each sampling occasion (Fig. 1). The number of leeches collected was negatively correlated with the Keetch–Byram drought index (climate data from the Deramakot Forestry Office, see Fig. S5), which increased during our study and corresponds to a decrease in the amount of rainfall in the area between the sampling occasions. It is also conceivable that each collection might have depleted a local leech population, and they might not have been mobile enough to replenish the sampling quadrat before the next collection. Based on our data and the current knowledge about the ecology of leeches, we suggest however to collect leeches during the rainy season. This could raise logistical obstacles and make it impossible to carry out concurrent camera trapping and leech sampling, since camera trapping is often carried out in the dry season. The two methods, however, could also complement each other, allowing surveying the mammal community throughout the year. This combined dataset could be analysed in a multi-season occupancy framework.

In contrast to Weiskopf *et al*. (2018), our data did not support the hypothesis that leeches are more suitable than camera traps to survey smaller mammal species. We detected no rats and mice, and although we detected one frog species in the leeches, our data showed a bias of leech detections towards larger bodied species, particularly ungulates. In fact, the proportion of ungulate detections in relation to detections of other mammals was higher in leeches than in camera traps (Fig. 3). In spite of this apparent bias, we also detected several other mammal species using the leeches, such as the critically endangered Bornean orangutan, the critically endangered Sunda pangolin, and the vulnerable and primarily arboreal binturong (a species not recorded by the camera traps).

Our analysis did reveal a difference in mammal detection success between the two types of leeches. It is possible that the larger size of tiger leeches leads to a higher chance to amplify mammalian DNA (see also Weiskopf *et al.*, 2018). We also found a slightly stronger bias in tiger leeches towards larger bodied mammals. This might be a result of their ecology, as tiger leeches live in small trees and bushes about 1 m off the ground (Lai *et al.*, 2011), while brown leeches mainly occur on the ground (Fogden and Proctor, 1985). Using an occupancy analysis where we estimated different detection probabilities for the two types of leeches allowed us to account for these differences. We are, however, aware that this might be challenging in other studies, due to the poorly known leech taxonomy (Schnell *et al.*, 2015) and the difficulties in distinguishing leech species in the field (Weiskopf *et al.*, 2018). Overall, average detection probability of the two leech types was lower than that of the camera traps, but for a number of species, tiger leeches had higher detection probabilities than camera traps (Fig. 4). As very low detection probabilities can result in poor occupancy estimates (MacKenzie *et al.*, 2002), future leech studies might be restricted to species with higher detection probabilities, or be able to increase sampling effort. This also highlights the need to account for varying detectability; conservation scientists should avoid using the raw number of detections as any kind of relative abundance measure (see Sollmann *et al*. (2013) for a discussion of the use of relative abundance index).

Despite the above differences, we found consistent estimates of occupancy probability across our camera-trap and leech datasets. This indicates that the leech method provides reasonable estimates of occupancy and is thus a suitable method for studying and monitoring mammal species in tropical rainforests. The occupancy estimates from the camera-trap data, however, had narrower BCIs, which was likely a result of the overall larger dataset. The smallest BCIs and likely the most robust and accurate measure of occupancy were derived by combining the leech and camera-trap datasets, due to the increased total amount of data available. Similarly, other occupancy studies that have used multiple detection methods reported improved occupancy estimates (Iknayan *et al.*, 2014; Nichols *et al*., 2008). The use of two detection methods allows researchers to collect more data, which will be especially beneficial in situations where detection probability is low and when rare species are the target. In camera-trapping studies, detection probability often depends on the way the camera traps are set up. Certain species such as larger felids are known to travel on roads or trails, whereas other species often avoid such features (Wearn *et al*., 2013). On the other hand, according to our results, the leech iDNA method preferentially samples larger ungulates. A key point for practitioners trying to choose between these two methods is to consider which method is more efficient for sampling. Camera trapping might generate more detections, since cameras can be left in the field for months, but iDNA benefits from the ease of leech collection (i.e. with the help of local people). In particular, iDNA could be the only feasible method for gathering large numbers of vertebrate detections during one-off visits to remote sites. One-off visits can yield data suitable for occupancy modeling if each sampling unit can be subdivided into several independent spatial replicates, which take the place of the typical temporal repeat visits (Guillera-Arroita, 2011). This, however, requires that if a species is present in one spatial replicate, it is present in all replicates - the spatial analog to the closure assumption, which is likely to be violated when habitat is not homogeneous.

In this study, we were mainly interested in examining whether the analysis of leech-iDNA could be used as an efficient tool for biodiversity assessments. Our nested study design most likely resulted in spatial autocorrelation between observations of neighboring stations for some species, particularly for wide ranging species such as the banteng or sun bears. This did not matter for our purpose of comparing the two types of data. Future studies that aim to apply the leech method for biodiversity assessment should take spatial autocorrelation into account.

We also acknowledge that leeches themselves move, and thus, our sampling locations might not coincide exactly with vertebrate presence locations. Although precise ecological information about the movement of terrestrial leeches is unavailable, leeches are believed to be mostly quiescent between feeding events (Schnell *et al.*, 2018). The consistency in occupancy modelling results from camera traps and leeches suggests that the potential movement of leeches does not cause significant bias in our study, but we note that we did not include any habitat covariates in our occupancy analysis. Leech movement could, however, cause potential problems when exploring species-habitat relationships, particularly at a fine scale in heterogeneous habitats. This problem increases for flying invertebrates, such as mosquitos, tsetse flies or carrion flies (Calvignac-Spencer *et al.*, 2013a), that likely move over larger distances of up to a few kilometers (Verdonschot & Besse-Lototskaya, 2014), or for eDNA samples which are regularly transported away from their original deposition sites to their collection sites by currents or wind. In concrete terms, a vertebrate obviously must be in front of the camera at the moment its photo is taken. With e/iDNA, the vertebrate does not have to be at the collection location, but could have been at any distance that the sample has moved since deposition. This could lead to wrong inferences about species habitat preferences. For this reason leeches, as well as ticks, present an advantage over other, more mobile invertebrates or samples taken from streams and rivers. Further real-world complications with e/iDNA that must be considered in future studies are, for example, different habitat and/or feeding preferences of invertebrates that affect species detection probabilities. On the other hand, the use of e/iDNA can increase sampling efficiency, making it possible to gather more detection events per unit effort. Proper sampling design and statistical modelling must therefore be used to correct for the extra uncertainty introduced by the use of e/iDNA samples so that the efficiency benefit of e/iDNA can be properly exploited. In this study we presented examples how occupancy models can help to overcome some of these real-world complications and we discuss opportunities how our approach could be extended by applying multi-season occupancy models or by using spatial replicates, for example.

Our results are a promising indication that use of iDNA can help to overcome difficulties in surveying and monitoring terrestrial mammals in tropical rainforests. The species accumulation curves and occupancy estimates indicate that the leech iDNA method performed similarly to the well-established camera-trap approach. The iDNA approach, however, was limited by sample sizes, and it may be challenging to collect sufficient samples to achieve accuracy in estimates comparable to that from camera-trapping. This suggests that the collection of iDNA may be best used to supplement camera-trap surveys. Leeches helped to detect a few species that were not detected during the camera-trap survey; allowed us to distinguish between similar species that could not be differentiated in photographs; and combining leech and camera-trap data in a single model improved estimates of occupancy estimates. The main challenge for upcoming studies relying solely on iDNA therefore appears to be the collection of a sufficient number of leeches, which may be helped by the use of invertebrate traps. In conclusion, iDNA presents a promising approach for systematically surveying wildlife populations, but future studies need to consider (a) potential sample size limitations and (b) idiosyncrasies of the detection data, such as the potential mismatch of detection and presence location, or factors influencing detection probability. In combining systematic leech surveys with occupancy modeling while accounting for differences in detection due to leech type, numbers, and detections of other species, we hope to highlight this approach to wildlife ecologists as a new sampling tool, and to molecular ecologists as a robust analytical framework for e/iDNA.

## Acknowledgements

We thank the Sabah Biodiversity Center and the Sabah Forestry Department, especially Johnny Kissing and Peter Lagan for support and involvement in this project. Many thanks go to the field team for their hard work while conducting the leech and camera-trap surveys. We thank Sebastian Wieser, Riddhi Patel, Anke Schmidt, Renata Martins, and Dorina Lenz for their technical advice on laboratory issues and sequencing work. This project received financial support from the German Federal Ministry of Education and Research (BMBF FKZ: 01LN1301A), Point Defiance Zoo and Aquarium through Dr. Holly Reed Conservation Fund and San Francisco Zoo.

## Data accessibility

Scripts are available in the supporting information of this article. The data supporting this article is not publicly available as it contains sensitive information about endangered and threatened species. For requests for use of the data for academic purposes please contact.

## Author contributions

AW and RS designed the study; AM and STW performed the sampling; JFA, LH, JA, ACP, RB, and DWY analyzed data; JFA and AW lead writing the manuscript. All authors contributed to drafts and gave final approval for publication.

